# Rapid resistance evolution against phage cocktails

**DOI:** 10.1101/2024.08.27.609872

**Authors:** Baltus A van der Steen, Matti Gralka, Yuval Mulla

## Abstract

When bacteria are treated with multiple antibiotics simultaneously, resistance is highly unlikely to evolve. In stark contrast, resistance against multiple phages frequently arises during therapy. Why does resistance against multi-phage cocktails evolve so easily? Using a mathematical model, we show how the bacterial evolutionary dynamics and phage replicative dynamics uniquely intertwine, facilitating the rapid evolution of multi-phage resistance. As different phages replicate and become inhibitory at varying time points, bacteria can sequentially acquire resistance rather than simultaneously – increasing the chance of multi-resistance by orders of magnitude. Additionally, we predict and experimentally verify a regime where multi-phage resistance is robustly prevented. Our findings provide a framework for the rational design of phage cocktails to curtail resistance development. Resistance can be minimized by reducing the dose of the most potent phages or by using phages with longer latent periods, as this helps synchronize multi-phage selection.

---

Bacteriophages are viruses that obligately parasitize bacteria, providing a highly specific mechanism of action that spares the host’s commensal microbiota[1]. Although initially developed in the early 20th century and utilized extensively before the antibiotic era, phage therapy was largely sidelined in Western medicine following the mass production of broad-spectrum antibiotics[2]. The global rise of multidrug-resistant ‘superbugs’ has led to a clinical renaissance for phages, with recent retrospective studies reporting success rates as high as 77.2% in difficult-to-treat cases[4]. However, resistance to therapeutic phages often arises during treatment[4]. To delay or prevent resistance, therapy typically uses cocktails of multiple phages[5]. Classical evolutionary theory predicts that the chance of a resistant mutant decreases exponentially with the number of inhibitory compounds[6] underpins the success of multidrug therapies against diseases like HIV/AIDS[7], tuberculosis[8] and cancer[9]. For example, monotherapy for HIV/AIDS using either protease inhibitors or nucleoside analogues proved ineffective as resistance emerged rapidly[10]; however, their combination is highly effective because it is exponentially more difficult for the pathogen to simultaneously evolve resistance against multiple compounds with distinct mechanisms[11]. This exponential scaling of the chance of resistance with the number of compounds is typically implicitly[12] and sometimes explicitly[13–15] assumed to be true for phages as well - suggesting that resistance evolution should be near-impossible when multi-phage cocktails are used for therapy.

In stark contrast to this prediction, resistance against all components of a multi-phage cocktail is frequently observed during therapy[4, 12], suggesting that multi-phage resistance evolves much more rapidly than predicted by current evolutionary theory.

We show that this contradiction is resolved by accounting for a key difference between phages and conventional drugs: phages replicate, uniquely coupling population dynamics with resistance evolution. To study this combination of phage population dynamics and resistance evolution, we first develop a minimal model of phage therapy with multiple phages against a single pathogen. We show that the probability of resistance evolution hinges on whether two phages act roughly simultaneously to quickly eradicate the host, preventing resistance evolution before population collapse. We then confirm our model predictions experimentally using different sets of phages that either do or do not act simultaneously. Together, our results explain why multi-resistance emerges during phage therapy with cocktails and provides a framework for developing phage cocktails less prone to resistance evolution during treatment.

## Methods

### Model

To maximize the generality and tractability of our model, we use a deliberately minimalistic model of phage predation. Consistent with previous approaches[16, 17], we model strictly lytic phages adsorb to and burst bacteria, releasing several new phage particles, which can start subsequent infection cycles. Bacteria grow in a steady state and can spontaneously evolve resistance. The number of bacteria that is sensitive to all phages, *b*_0_, grows at a rate μ and spontaneously evolves resistance against phage *i*(*p*_*i*_) at a rate *r*_*i*_. We include predation by phage *i*, which adsorbs to bacteria at a rate *b*_0 *kadsorb, i*_:

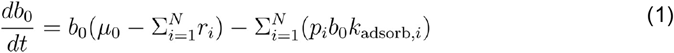

Next, we model the population dynamics of bacteria resistant to phage *j*(*b*_*j*_) and sensitive to all other phages. For simplicity, we do not consider resistance tradeoffs or phage (co-) evolution:

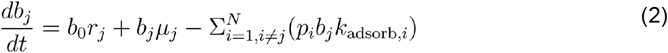

As our model is continuous and mutants resistant to both phage *j* and *l* can have very small population sizes, we only allow for cell doublings if there is at least one double-resistant mutant:

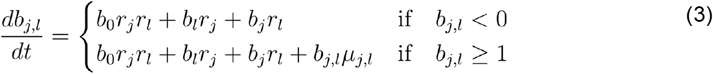

Bacteria infected by phage *j*(*I*_*j*_) burst after a latent period ofτ _*latency:*_

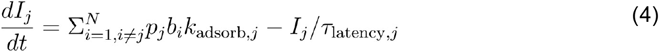

For each burst,*n*_burst_ phages are released. As bacteria typically evolve resistance via receptor protein mutations, we assume phages do not adsorb to resistant bacteria:

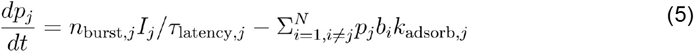

Lastly, we count the number of independently evolved dual-resistant bacterial mutants independently evolved dual-resistant mutants that have arisen in the population (so neglecting any subsequent divisions):

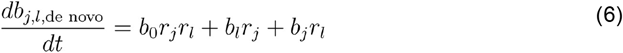

Next, we approximate the chance that a double-mutant evolves, 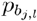. To this end, we calculate the chance of at least one mutation occurring, which follows a Poisson distribution according to the classical result of Luria and Delbruck[18]:

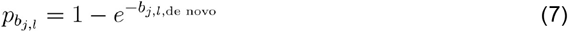

With this approach, we can approximate the chance of resistance despite the deterministic nature of the model: solving Eq. 6 yields the average number of double-resistant mutants *b*_*j,l*_ and by assuming that the number of double-resistant mutants is exponentially distributed over replicate experiments we can then calculate the chance of at least one double-resistant mutant per experiment using Eq. 7.

The set of differential equations was solved numerically using solve_ivp (scipy 1.11.1, Python 3.11.5) and the collapse time of the bacterial populations was detected using find_peaks (scipy 1.11.1, Python 3.11.5)[19].

### Chance of simultaneous selection pressure in a phage cocktail

Our model shows that the likelihood of resistance emerging depends critically on whether two phages exert selection pressures simultaneously or sequentially—a dynamic governed by their kinetic properties and initial concentrations. As more phages are added to a cocktail, the probability increases that at least one pair will exert simultaneous selection pressure.

To quantify this, we estimated the probability that a randomly assembled cocktail of *n* phages from a biobank of *N* phages contains at least one pair exerting simultaneous selection pressure. Assuming the number of such synergistic pairs is small relative to the total number of combinations(*N* (*N* − 1) / 2, the probability can be approximated :

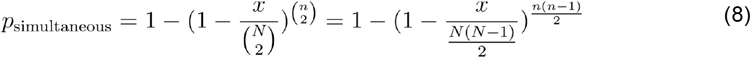

The value 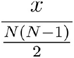 depends on the collapse time distribution in the biobank. We calculated this parameter using a previously published dataset[20] that measured collapse times for 69 phages from the publicly available Basel collection^11^, all infecting the same laboratory strain of *E. coli* (MG1655) under fixed inoculum conditions. Although this strain is non-pathogenic and the data is from in vitro experiments, to our knowledge it is the only available dataset with collapse times measured consistently across a large number of phages.

Suppl. Fig. 3a shows the distribution of collapse times for the individual phages, and Suppl. Fig. 3b–c illustrate the pairwise differences in collapse times. Assuming a 1-hour window of opportunity to prevent resistance (based on Suppl. Fig. 2), approximately 20% of phage pairs fall within this window (Suppl. Fig. 3). In Fig. 5, we plot Eq. 8 using 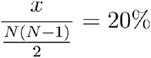, corresponding to this empirical estimate for the Basel collection, to show how the probability of simultaneous selection pressure scales with cocktail size.

### Bacteria and phage propagation

All experiments were performed using *Escherichia coli* MG1655 carrying the kanamycin-resistant plasmid pCS-λ, which encodes a luciferase-based bioluminescence reporter[21]. Pre-cultures were grown overnight at 30 °C in high salt lysogeny broth (LB) with shaking at 150 rpm for liquid cultures, or on 1.5% LB agar (LBA) plates for solid cultures. Phages p17, p33 and p61 of the Basel collection[22] were propagated on *E. coli* in LB at 30 °C with shaking (150 rpm) overnight. Phages were purified by chloroform treatment (1:75, v/v)[23], followed by centrifugation at 2800 × g. Phage concentrations were determined by 10-fold serial dilution in LB, followed by spotting of 3 μL droplets onto double-layer LBA plates (0.5% top agar) seeded with *E. coli*.

### Resistance frequency assay

Phage resistance frequencies were assessed using double-layer LBA plates. Phage-containing supernatants were mixed with overnight *E. coli* cultures and overlaid onto the bottom LBA layer to test resistance against p17, p33, p61, and their combination. To determine the bacterial inoculum concentration, a 10-fold serial dilution of the *E. coli* culture was plated on LBA and incubated overnight at 30 °C. Colony-forming units (CFUs) were counted to calculate resistance frequencies for each condition[18].

### One-step growth curve assay

*E. coli* was cultured to mid-log phase (OD_600_ ≈ 0.5), diluted (1:100, v/v) and infected with either phage p17, p33 or p61 (final concentration: 10^5^ PFU). After a 5-minute adsorption period, cultures were again diluted 1:100 (v/v) and incubated at 30 °C. Samples were collected every 5 minutes, treated with chloroform (1:75, v/v), and phage concentrations were determined via 10-fold serial dilution and spotting onto double-layer agar plates seeded with *E. coli*[24]. The latent period, τ_lantent_, was estimated by empirically fitting the time-resolved phage concentration [*p*] with a Hill function 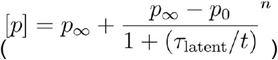 using optimize.curve_fit (SciPy 1.13.1) in Python[19].

### Single phage growth curve

To quantify phage-induced collapse times at various concentrations, 5-fold serial dilutions of either p17 (3 × 10^9^ PFU/mL), p33 (3 × 10^10^ PFU/mL) and p61 (6 × 10^9^ PFU/mL) were prepared. *E. coli* was grown to mid-log phase, diluted to an OD_600_ of 0.05 (1.69 × 10^7^ CFU/mL), and infected with each phage dilution (1:200, v/v). Cultures were incubated in a 96-well plate (200 μl/well) at 30 ^0^C for 48h with double orbital shaking. OD_600_ and bioluminescence (BioTek Synergy HTX Multi Mode Plate Reader) were recorded every 10 minutes to determine the collapse time and any regrowth. Collapse times were extracted from bioluminescence curves using find_peaks (SciPy 1.1.13.1) with a smoothing width of three data points.

### Dual-phage cocktail

Phages p17, p33 and p61 were diluted to match the concentrations used in the 5-fold serial dilution series from the individual phage growth curve experiments, resulting in an inoculum concentration range of 10^3^-10^7^ pfu/ml. For comparison, phage preparations at the Belgian phage therapy center[4] are administered at 10^6^-10^7^ pfu/ml. However, the effective inoculum concentration at the site of an infection is likely much lower than that of the preparation administered to a patient. The pharmacokinetics of phage products are still poorly understood and likely differ strongly per type of infection. Therefore, although we use realistic phage cocktail concentrations as a guide, we believe it is not yet possible to quantitatively compare our results to actual phage therapy due to a lack of pharmacokinetic knowledge. *E. coli* was grown to mid-log phase and diluted to an OD_600_ of 0.05. Phages were then combined in a 96-well plate (200 μl/well) in a two-dimensional gradient (checkerboard assay)[25], testing all pairwise combinations of the two dilution series across a 48-hour incubation period with shaking at 30 °C. OD_600_ measurements were taken at 0 h for background subtraction and at 48 h to assess bacterial growth. The background-subtracted OD_600_ values were plotted against the difference in collapse time, as determined from the corresponding single-phage growth curves.

## Results and Discussion

Using a minimal mathematical model, we first model a bacterial population exposed to a single type of phage (Methods, Suppl. Table 1, reference [26]). We observe that the bacterial population initially grows seemingly unperturbed by the presence of the phage until, after several hours, the bacterial population suddenly collapses due to phage predation, followed by regrowth driven by resistant mutants (Fig. 1a). The bacterial growth is initially unaffected by the phage titer as most bacteria remain uninfected until phages have replicated to much higher numbers[27]. The timing of the collapse is accelerated as the phage inoculum concentration increases (red to blue), in accordance with empirical observations[20].

**Fig. 1.**
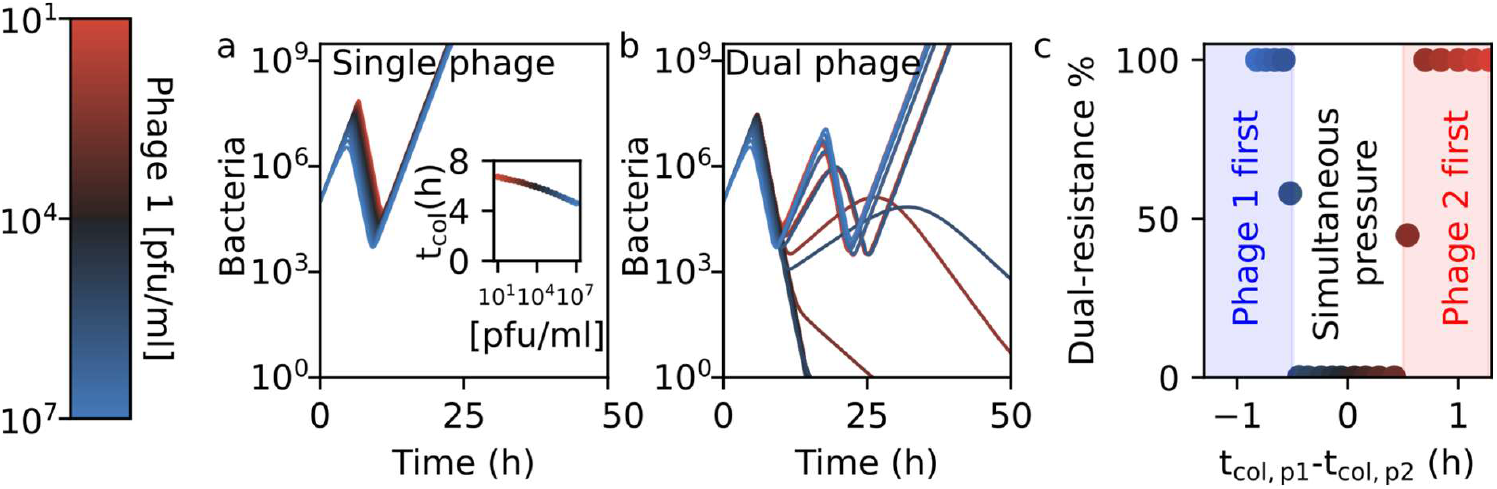
Theory predicts timing of selection pressure determines the likelihood of multi- phage resistance. **a)** Simulated bacterial population dynamics under attack from a single phage. Growth initially continues before a sudden collapse and rebound due to resistant mutants. Inset: collapse time as a function of phage inoculum concentration (red to blue, see color bar and inset). **b)** Dynamics under a two-phage cocktail, varying the inoculum concentration of phage 1 (same as in panel a, see color bar), keeping the concentration of phage 2 fixed at 10^4^ pfu/ml - with the black line representing equal inoculum concentrations for both phages. Depending on the inoculum concentration of phage 1, phage action can be simultaneous (single collapse in case of the dark lines) or sequential (two collapses in case of the bright red and bright blue lines). **c)** Probability of double-resistance emergence (calculated from simulations in panel b) as a function of the difference in collapse times between the individual phages 1 and 2 (calculated from panel a). Resistance is strongly suppressed only when both phages act simultaneously. All Panels share the same phage inoculum concentration color scale (left).

We investigate a bacterial population exposed to two distinct phages, differing (for now) only in their inoculum concentrations, and find two qualitatively different regimes. When both phages have similar inoculum concentrations, the population undergoes a single and definitive collapse (Fig. 1b, dark curves). In contrast, if inoculum concentrations differ substantially, we observe two sequential collapses followed by a definitive regrowth: the first due to one phage, the second following regrowth of a single-resistant mutant and subsequent collapse due to the second phage and regrowth of a dual-resistant mutant (Fig. 1b, blue and red curves). We call these two regimes simultaneous and sequential selection, respectively.

We calculate the chance of double-resistant mutants arising by first computing the expected average number of double-resistant mutants per experiment (Method Eq. 6) and then calculating the chance that at least one mutant arises per experiment. In the single-collapse regime (simultaneous selection), the probability of double-resistance emergence (Methods Eq. 7) is very low (<<1%, Fig. 1c) because the host population size declines too rapidly for double mutants to emerge. In contrast, in the double-collapse regime (sequential selection), the chance of escape mutants rapidly increases to near-certainty as double resistance can evolve in steps. Therefore, we conclude that the evolution of multi-phage resistance is only prevented within a window of opportunity to prevent resistance where both phages are selective simultaneously (.Fig. 4).

This window of opportunity to prevent resistance provides an experimentally testable prediction: resistance is prevented only when both phages have a matched collapse timing. We test this prediction using two *Escherichia coli* phages (p33 and p61 of the BASEL collection[22]) with similar collapse times (Supplementary Table 1, column 3 of Ref. [20]) but distinct receptor proteins (Supplementary Table 1, column 28 of Ref. [22]), suggesting different resistance pathways. When exposing the host with only one phage at a time, each phage yields resistance in ∼1 per million cells, but dual resistance is undetectable (<10^−9^, Suppl. Fig. 2a).

Previous work simulating bacterial growth curves[28] has revealed that the collapse times depend on both phage parameters (burst size, adsorption rate, latency time), but also initial phage abundance. Therefore, we systematically vary the inoculum concentrations of each phage across four orders of magnitude and measure bacterial growth after 48 hours. Single phage treatments always result in regrowth, and regrowth is also observed in ^⅓^ of dual-phage cocktails (Suppl. Fig. 1b) — despite the number of bacteria being much smaller than the inverse frequency of dual resistance (Suppl. Fig. 1a). Closer inspection reveals that successful suppression without regrowth only occurs at high inoculum concentrations of p61 but low inoculum concentrations of p33 (Fig. 2c).This means that an increase in p33’s inoculum concentration actually decreases suppression - counterintuitively, because typically, one might expect an increased dosage to lead to greater efficacy. More specifically, we find that resistance is only suppressed when the collapse time of p61 precedes that of p33 (Fig. 2d).

**Fig. 2.**
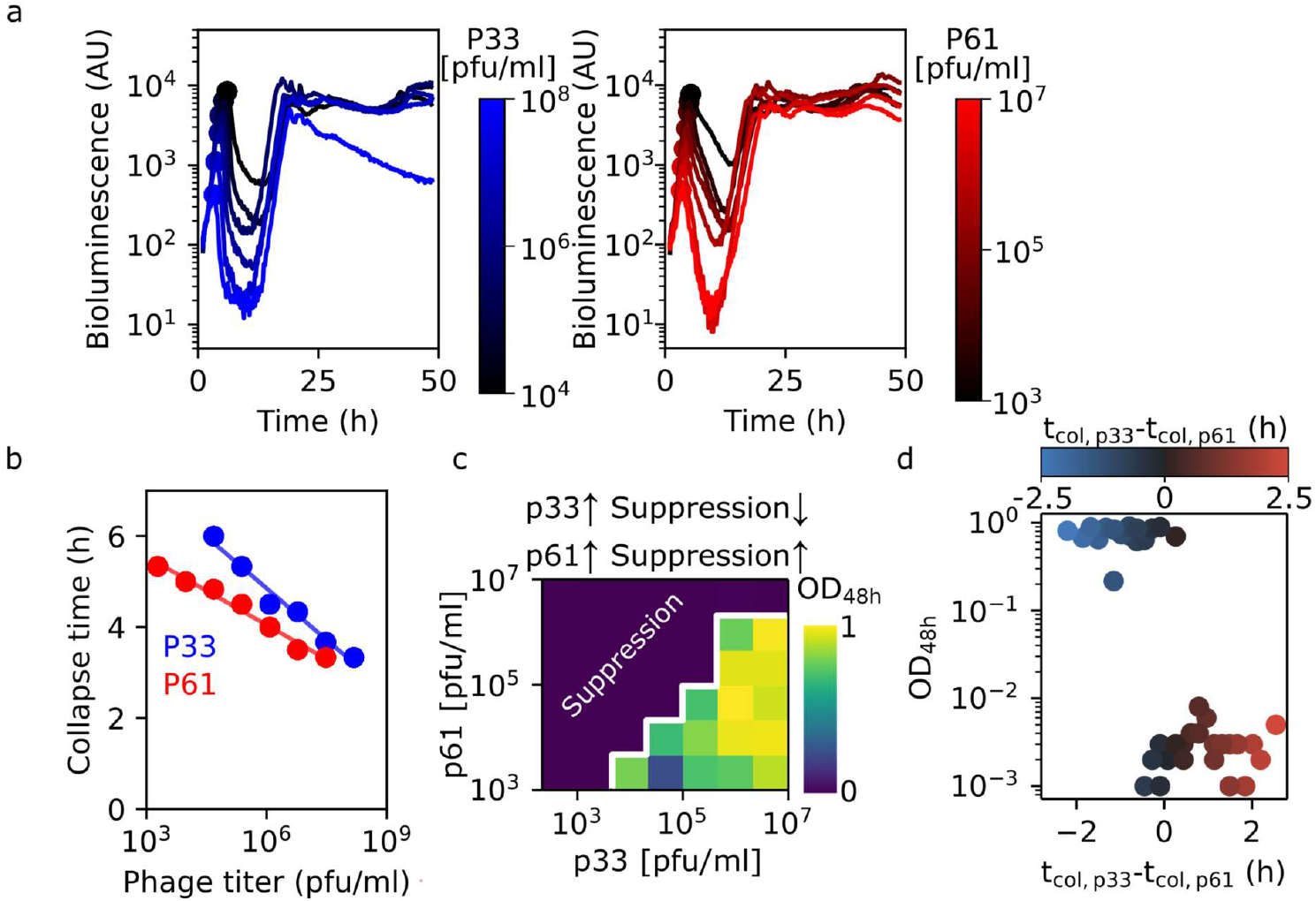
Experiments confirm timing of selection pressure determines the likelihood of multi-phage resistance. **a)** Bacterial growth curves at increasing phage concentrations: left panel for p33 (blue to black), right panel for p61 (red to black). **b)** Collapse times derived from growth curves in panel a. **c)** Experimental data showing bacterial growth (optical density after 48 h) is only suppressed when p61 is at high and p33 at low inoculum concentrations, meaning that increasing the inoculum concentration of p33 actually decreases suppression. The white line highlights the sharp transition where phages no longer stably suppress bacterial growth. **d)** The same data as panel c, now plotted against the difference in single-phage collapse time (panel b).

Similar to our model prediction (Fig. 1c), we experimentally find that the difference in collapse time is the main determinant for stable suppression of bacterial growth (Fig. 2d). However, in contrast to the predictions of our simple model (Fig. 1c), we do not experimentally observe a window of opportunity to prevent resistance that is symmetric around 0 difference in collapse time. However, that model made the strong assumption that phage parameters were identical between the phages. Lifting this assumption, we find that the window of opportunity to prevent resistance predictably depends on these phage parameters: increases with the phage latent period widen the window (Suppl. Fig. 2) and asymmetry arises when phages have different latent periods. Therefore, resistance is best prevented when the phage with the longer latent period acts first (Fig. 3a). After determining the latent period of p33 and p61 via one-step growth curve experiments, we find that, consistent with our theoretical prediction, p61 (the phage which suppresses regrowth when causing the early collapse) indeed has a longer latent period than p33 (resp. 49±3 min and 37±1 min, p=10^−5^, Fig. 3b, Suppl. Fig. 1c).

**Fig. 3.**
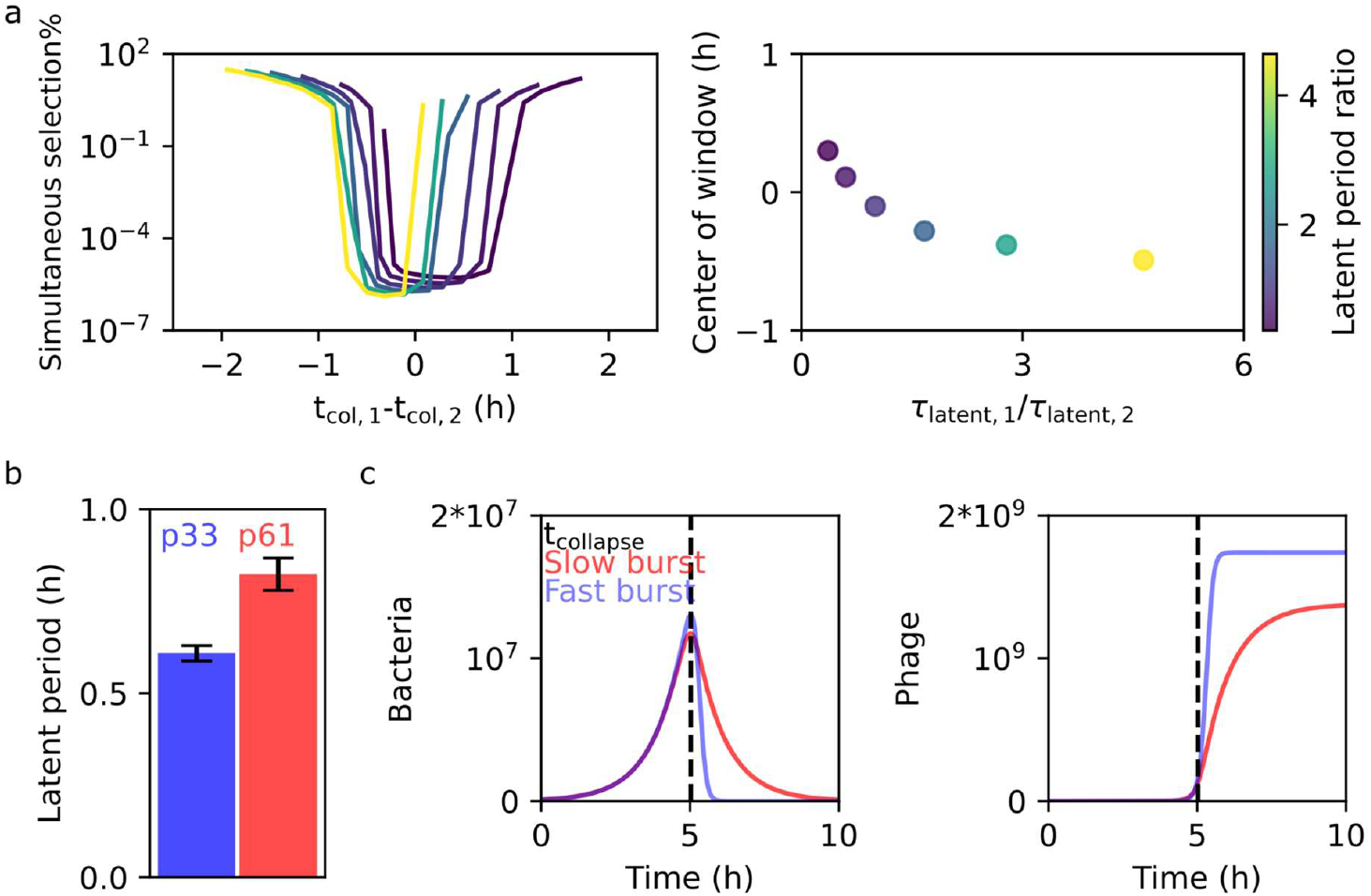
Resistance is suppressed most when the phage with the longest latent period causes the first collapse. **a)** Dual-phage simulations with identical burst size and adsorption rate, differing only in their latent period (color bar, see suppl. Table 1 for all parameters). We vary the difference in collapse time (x-axis, left panel) by varying the inoculum concentrations. The theoretical model predicts that the window where resistance is suppressed shifts as a function of the ratio between the latent periods of both phages (right panel), such that resistance is suppressed most when the phage with the longest latent period causes the first collapse. **b)** We experimentally confirm that phage 61, which has to collapse first for efficient resistance suppression (Fig. 1c), indeed has a longer collapse time. The error bar shows the standard deviation of the fit (Suppl. Fig. 1c). **c)** Simulations of single-phage predation assays reveal why the latent period shifts the window of opportunity to prevent resistance. If the latent period is short (blue), then the collapse of the bacterial population (right) and the increase in the phage population (left) are sudden. In contrast, both are more gradual if the latent period is long (red) - shifting the window a second phage can simultaneously suppress growth.

To understand the mechanism by which the latent period affects the window of opportunity to prevent resistance, we perform simulations of bacteria exposed to a single type of phage, varying the latent period of that phage. These simulations reveal why the latent period is such a relevant parameter: whereas a short latent period causes a steep collapse, a long latent period causes a more gradual collapse (Fig. 3c). Such a gradual collapse broadens the window of opportunity to prevent resistance and allows the second phage to catch up with the phage causing the first collapse.

In our experiment, p33 and p61 were carefully chosen for their similar bactericidal activity (comparable collapse times). Indeed, when we choose sets of phages with very dissimilar bactericidal activity (p17 and p61), the robust resistance suppression is only achieved at an extreme inoculum concentration difference (>100x, Suppl. Fig. 4). This shows that, following the prediction, collapse times roughly must coincide for resistance suppression, raising the question how this can be achieved when assembling phage cocktails. To highlight how likely an arbitrary cocktail of phages is to operate within this effective window as a function of the number of phages in a cocktail, we analyzed published collapse-time data from multiple phages infecting *E. coli*[20]. We calculate the likelihood that any two phages within a random cocktail fall within this window (Methods Eq. 7), leading to simultaneous selection. We find that this chance increases roughly linearly for low numbers of phages (Fig. 5), which is qualitatively different from the exponential scaling of chance of resistance for antibiotics^4^. This qualitative difference between antibiotics and phages as bactericidal agents may explain why multi-resistance is more frequently observed during phage therapy than multi-resistance against antibiotic cocktails.

**Fig. 4.**
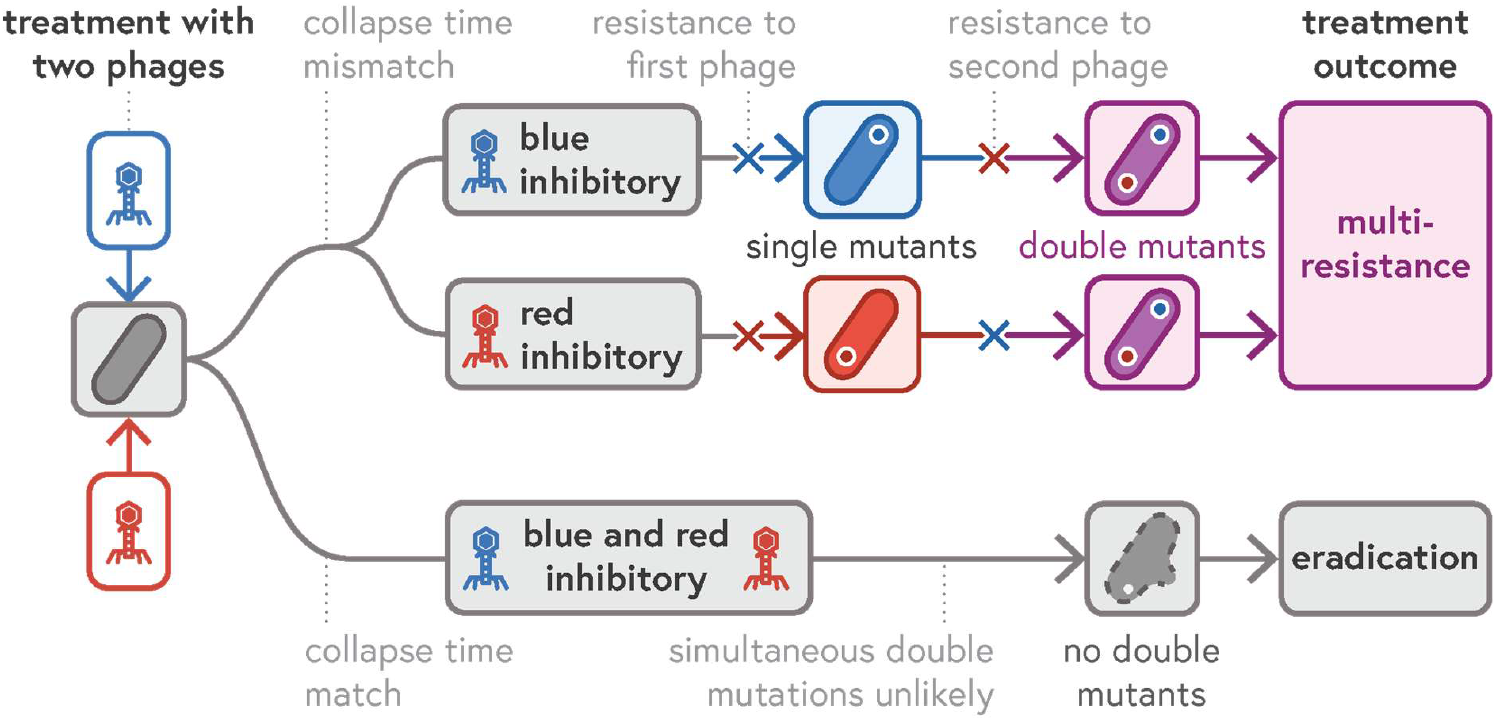
Schematic of the resistance evolution pathways against phage cocktails. Upon application of a multi-phage cocktail, each phage replicates at its own rate and reaches inhibitory concentrations at different times. If these selective events occur sequentially, resistance can evolve stepwise. Only when multiple phages act simultaneously is the evolution of multi-phage resistance robustly suppressed (bottom).

**Fig. 5.**
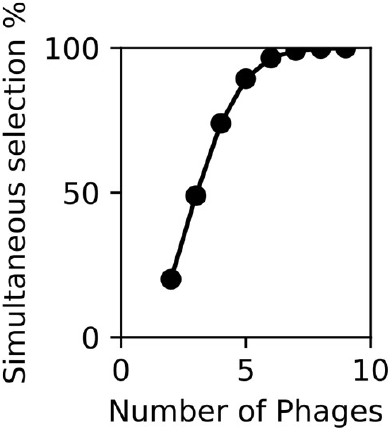
Complex enough phage cocktails still robustly suppress resistance evolution. Simulated probability that at least one phage pair in a cocktail exerts simultaneous selection, plotted as a function of cocktail size and phage latent period. Based on data from a public *E. coli* phage library[20, 22]. Calculations follow Methods Eq. 8. The plot shows that even though multi-phage cocktails are less effective than traditional drugs in suppressing resistance, simultaneous selection pressure (which effectively prevents resistance) can still be achieved by combining enough different phages.

Our study explains how multi-phage resistance evolution may differ from that of antibiotics due to the replicative nature of phages and offers an explanation of why multi-phage resistance is a major clinical problem. Although we have only considered de novo mutations, resistance acquired via horizontal gene transfer would follow similar rules as the only alteration would be that the mutation rate then also depends on the density of donor bacteria as reviewed in Ref. [30]. Therefore, multi-phage resistance can be acquired sequentially independent of the mode by which the resistance evolves.

In principle, *in vitro* screening could help identify phage pairs that cause similar collapse timings. However, phage pharmacokinetics *in vivo* differ markedly from test tube conditions[31], so predicting collapse timing in patients is difficult. This discrepancy suggests that in vitro cocktail performance should be interpreted with caution; a cocktail’s apparent efficacy in a test tube may rely on a specific timing of selection pressure that does not translate to the patient. Our work does show that the initial phage titer controls evolutionary dynamics in non-trivial ways. Therefore, an important next step is to record resistance rates as a function of initial phage titers in more complex infection models (e.g. animal models[32] or organoids[33]) to find optimal regimes.

Designing effective phage cocktails requires a rigorous understanding of the evolutionary interplay between constituent phages. For example, resistance to one phage must not confer cross-resistance to others in the mixture[34]. Our findings demonstrate that standard *in vitro* cocktail assays can conflate these fundamental evolutionary interactions with population dynamics, such as collapse timing. Because these timing effects are highly sensitive to pharmacokinetic variations and likely translate poorly to *in vivo* conditions, it is critical to employ diagnostic assays that isolate evolutionary outcomes from ecological noise. Researchers can directly quantify the genetic barriers to multi-phage resistance through sequential exposure, which involves isolating mutants resistant to a primary phage and then challenging them with a secondary agent. Unlike simultaneous cocktail assays[35, 36], these sequential tests[37] are independent of collapse timing and thus provide a more robust assessment of a cocktail’s therapeutic potential in clinical settings. Sequential testing reveals cross-resistance patterns[38], but does not ensure a simultaneous selection pressure. To increase the change of simultaneous selection and therefore robust resistance suppression, it is beneficial to further increase the number of distinct phages (Fig. 5) - especially those with longer latent periods (Suppl. Fig. 2b). Finally, resistance does not always imply failure[4, 39]: resistance to phages can increase sensitivity to antibiotics[40] or reduce virulence[41]. These evolutionary trade-offs could be exploited, and some researchers already recommend including them as a criterion in phage selection[42]. Our findings highlight the challenge of fully preventing phage resistance, the importance of better understanding *in vivo* phage pharmacokinetics and the need to integrate evolutionary principles into therapy design.

---

**Suppl. Fig. 1.**
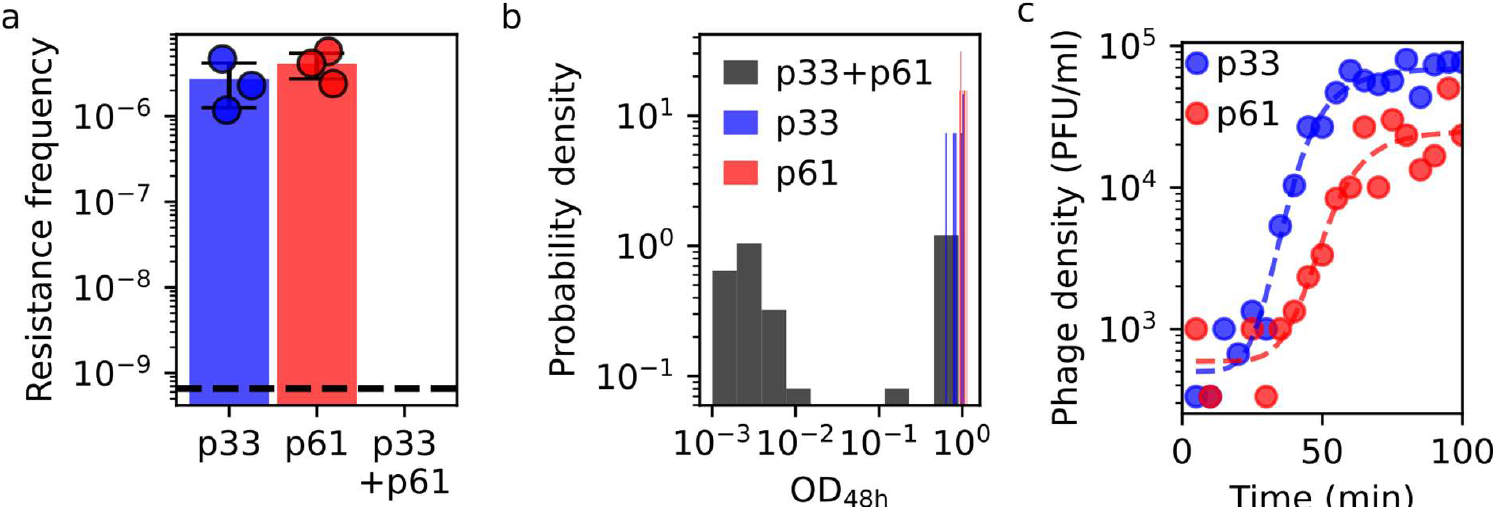
p33 and p61 characterization. **a)** Resistance frequencies for p33 alone (blue), p61 alone (red), and both combined (below detection limit; dashed line). **b)** Final optical density distribution after 48 h for p33 (blue), p61 (red), and the cocktail (grey). **c)** One-step growth curves for p33 and p61.

**Suppl. Fig. 2.**
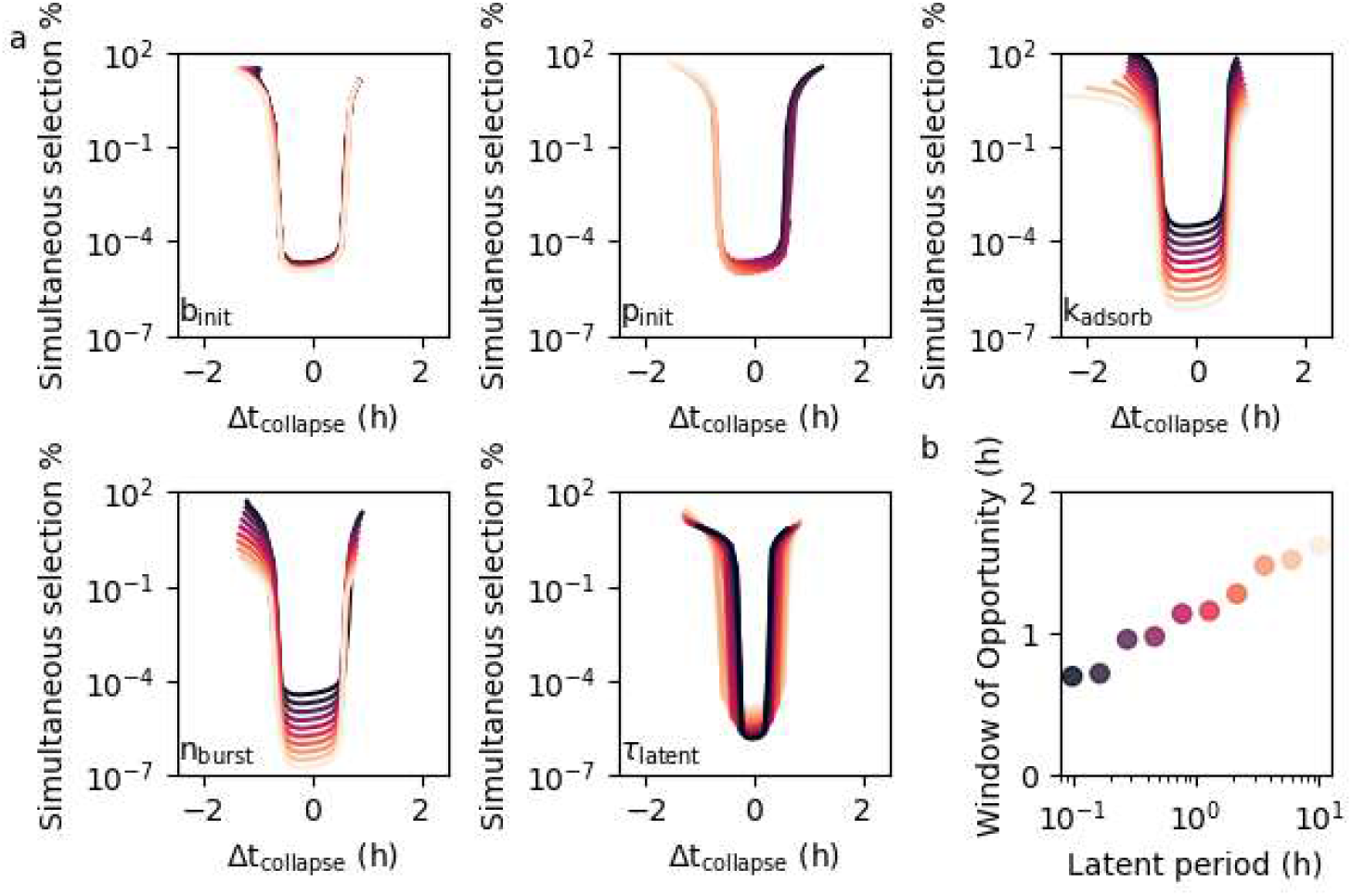
Latent period influences the size and position of the window of opportunity to prevent resistance. Dual-phage simulations using identical phages, varying the inoculum concentration of one phage holding the other constant. Resistance frequency (y-axis) is plotted against the difference in collapse time (x-axis) derived from single-phage simulations. Simulations were run across a range of parameters (different panels) in logarithmic steps (ranging from low in dark to high in bright, see Suppl. Table 1 for the exact ranges). **b)** Wider latent periods increase the window of collapse-time differences over which resistance is suppressed. The color coding reflects the latent period and is the same as in panel a, bottom-right.

**Suppl. Fig. 3.**
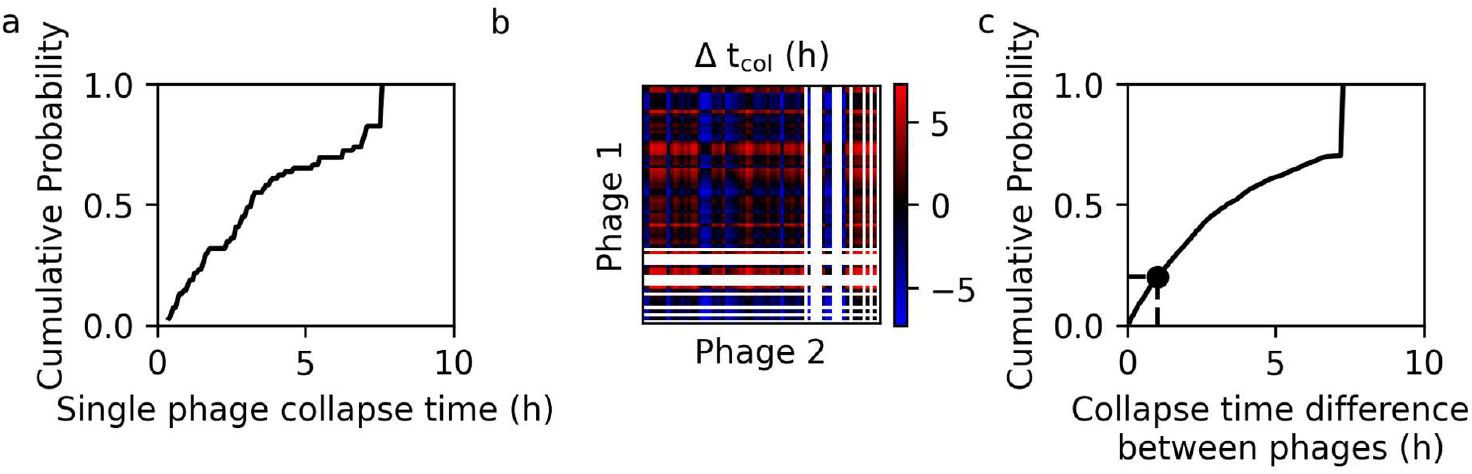
BASEL collection collapse time distribution. To estimate how often two phages might act simultaneously and thus prevent resistance, we analyzed published collapse-time data for 69 *E. coli*-infecting phages from the BASEL collection[20]. **a)** Cumulative distribution of collapse times. For 11 of 69 phages, collapse times could not be measured due to very slow lysis dynamics. **b)** Heatmap showing pairwise differences in collapse time for all measurable phage pairs. White lines indicate phages for which collapse times were undetermined. Collapse time differences between specific phage pairs can be found in the Supplementary Data. **c)** Cumulative distribution of collapse-time differences from panel b, excluding the diagonal (self-comparisons). Pairs involving phages with indeterminable collapse times were treated as having infinite differences. ∼20% of phage-phage pairs differ by 1 hour or less (dashed line).

**Suppl. Fig. 4.**
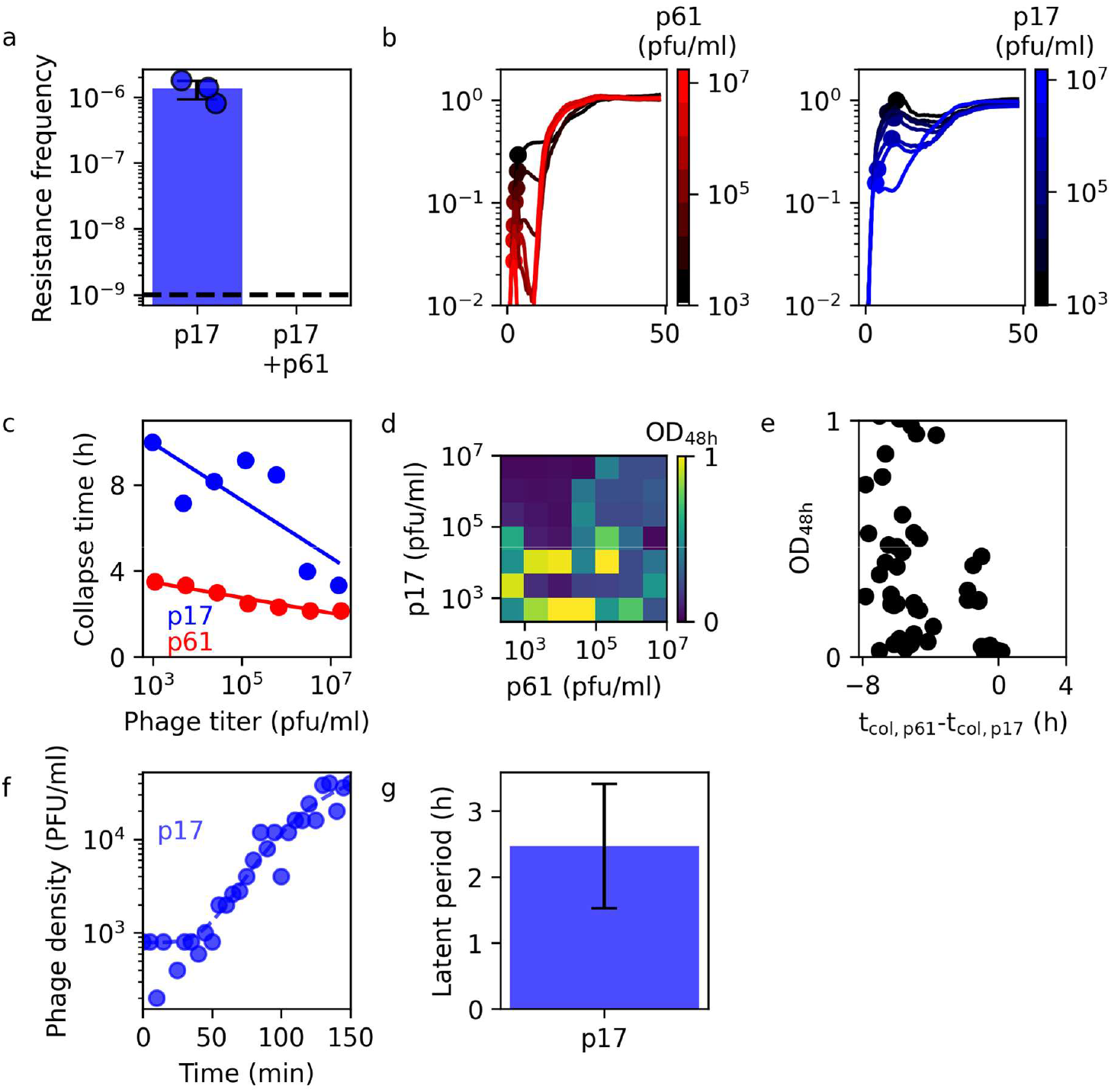
phages with different bactericidal activity only show stable suppression at a large phage inoculum concentration imbalance. **a)** Resistance frequencies for phage 17 alone (blue) or combined with p61 (below detection limit; dashed line, see p61 alone is in Suppl. Fig 1a). The results indicate no collateral resistance between both phages. **b)** Bacterial growth curves at increasing phage concentrations: left panel for p17 (blue to black), right panel for p61 (red to black). **c)** Collapse times derived from growth curves in panel b. **d)** Experimental data showing bacterial growth (optical density after 48 h) is only suppressed when the inoculum concentration of p17 is at least 100x higher than of p61. **e)** The same data from panel c, now plotted against the difference in single-phage collapse time (panel b). **f)** One-step growth curves for p17. **g)** Latent period extracted from the one-step growth curve (panel f). The error bar shows the standard deviation based on the fit.

**Suppl. Table 1.** .Model parameters used in simulations. To explore the impact of different phage traits on resistance evolution, we simulated a dual-phage predation assay in which both phages initially share identical kinetic parameters. Each parameter was varied individually across 10 logarithmically spaced steps, keeping all others constant. To modulate the difference in collapse timing, the inoculum concentration of one phage was varied across 20 logarithmically spaced steps from 10^1^ to 10^7^ pfu, keeping the concentration of the other fixed at 10^4^ pfu.

| | $b_{\text{init}}$<br>[cfu] | $p_{\text{init}}$<br>[pfu] | $\mu$<br>[h<sup>-1</sup>] | $k_{\text{adsorb}}$<br>[h<sup>-1</sup> cfu<sup>-1</sup>] | $\tau_{\text{latency}}$<br>[h] | $n_{\text{burst}}$<br>[ ] | $r$<br>[ ] |
| --- | --- | --- | --- | --- | --- | --- | --- |
| Fig. 1 | 10<sup>5</sup> | 10<sup>1</sup>-10<sup>7</sup> | 1 | 10<sup>-8</sup> | 1 | 100 | 10<sup>-6</sup> |
| Suppl. Fig. 3<br>(fixed) | 10<sup>5</sup> | 10<sup>1</sup>-10<sup>7</sup> | 1 | 10<sup>-8</sup> | 1 | 100 | 10<sup>-8</sup> |
| Fig. 3 a, Suppl.<br>Fig. 3<br>(variable) | 10<sup>4</sup>-10<sup>6</sup> | 10<sup>3</sup>-10<sup>5</sup> | 0.2-2 | 10<sup>-7</sup>-10<sup>-9</sup> | 0.02-2 | 50-5000 | 10<sup>-8</sup> |

